# Integrative analysis of single cell expression data reveals distinct regulatory states in bidirectional promoters

**DOI:** 10.1101/315937

**Authors:** Fatemeh Behjati Ardakani, Kathrin Kattler, Karl Nordström, Nina Gasparoni, Gilles Gasparoni, Sarah Fuchs, Anupam Sinha, Matthias Barann, Peter Ebert, Jonas Fischer, Barbara Hutter, Gideon Zipprich, Bärbel Felder, Jürgen Eils, Benedikt Brors, Thomas Lengauer, Thomas Manke, Philip Rosenstiel, Jörn Walter, Marcel H. Schulz

**Affiliations:** Excellence Cluster for Multimodal Computing and Interaction, Saarland Informatics Campus, Saarland University, Saarbrücken Germany; Department of Computational Biology and Applied Algorithmics, Max Planck Institute for Informatics, Saarland Informatics Campus, Saarbrücken, Germany; Graduate School of Computer Science, Saarland University, Saarbrücken, Germany; Department of Genetics, University of Saarland, Saarbrücken, Germany; Max Planck Institute of Immunobiology and Epigenetics, Stübeweg 51, Freiburg, 79108, Germany; Institute of Clinical Molecular Biology, Christian-Albrechts-University, Kiel, Germany; Applied Bioinformatics, Deutsches Krebsforschungszentrum, Heidelberg, Germany; Data Management and Genomics IT, Deutsches Krebsforschungszentrum, Heidelberg, Germany

**Keywords:** bidirectional genes, single cell RNA-seq, epigenetics

## Abstract

**Background:** Bidirectional promoters (BPs) are prevalent in eukaryotic genomes. However, it is poorly understood how the cell integrates different epigenomic information, such as transcription factor (TF) binding and chromatin marks, to drive gene expression at BPs. Single cell sequencing technologies are revolutionizing the field of genome biology. Therefore, this study focuses on the integration of single cell RNA-seq data with bulk ChIP-seq and other epigenetics data, for which single cell technologies are not yet established, in the context of BPs.

**Results:** We performed integrative analyses of novel human single cell RNA-seq (scRNA-seq) data with bulk ChIP-seq and other epigenetics data. scRNA-seq data revealed distinct transcription states of BPs that were previously not recognized. We find associations between these transcription states to distinct patterns in structural gene features, DNA accessibility, histone modification, DNA methylation and TF binding profiles.

**Conclusions:** Our results suggest that a complex interplay of all of these elements is required to achieve BP-specific transcriptional output in this specialized promoter configuration. Further, our study implies that novel statistical methods can be developed to deconvolute masked subpopulations of cells measured with different bulk epigenomic assays using scRNA-seq data.

## 1 Background

Promoters are key structures for a coordinated regulation of gene expression. The increasing number of large-scale high resolution epigenomic and RNA-sequencing technologies are leading to a deeper understanding of genome-wide promoter configurations. Recent studies show that the number of bidirectional promoters (BPs) in the human genome is much larger than previously anticipated [1, 2, 3]. Sensitive assays, such as sequencing of nascent RNAs (GRO-seq) or 5^′^-ends of capped nascent RNAs (GRO-cap and Start-seq), allow the detection of unstable nascent RNAs produced at promoters, and have revealed more widespread bidirectional transcriptional initiation than previously recognized [4, 5, 6]. However, the exact classification of bidirectional or unidirectional promoters in a sample of interest is challenging, as it depends heavily on the sensitivity of the sequencing assay to recognize unstable, nascent RNAs [7, 8].

Recent studies discuss two types of bidirectional promoters. The first type concerns transcription of two RNAs in opposite direction from one core promoter, *i.e.,* one promoter leads to bidirectional transcription [9, 5, 10]. In the second type, transcriptional initiation of both RNAs occurs at two distinct core promoters that are close to each other, but are oriented in reverse direction, thus sometimes termed divergent bidirectional promoters. In this work we focus on bidirectional promoters that have two distinct core promoter elements that drive divergent transcription of two nearby genes.

BPs harbor overrepresented TF binding sites such as GABPA, MYC, YY1, NRF-1, E2F1 and E2F4 [11]. For example, the introduction of GABPA binding sites into unidirectional promoters lead to bidirectional expression in 67% of the cases [12]. Further, the sequence elements at some BPs function as inseparable units [13]. Other TFs prevent bidirectional expression, for example, promoters that show elongation in only one direction show enrichment of CTCF binding sites [4, 14]. However, more research is needed to investigate how TF binding determines directionality of initiation and elongation at BPs [9].

It was recently shown that the two Transcription Start Sites (TSSs) at a BP define a Nucleosome Free Region (NFR) between them. The size of the NFR may be an important structural element in BP regulation, determining the availability of binding sites for different TFs at the promoter and thus influencing gene expression strength as well as responsiveness to external stimuli [5, 6]. The current results point to a model, where an independent Pol2 complex assembles at each TSS and initiates transcription, such that accurate phasing of the +1 and −1 nucleosomes at these BPs allows epigenetic regulation through HMs [4, 5, 6]. Comparisons between BPs and unidirectional promoters suggest that HMs associated with active gene expression exhibit a bimodal distribution at BPs, and that upstream proximal enhancer marks may regulate downstream gene transcription [14, 6].

In summary, previous studies rely on the comparison of unidirectional against bidirectional promoters to learn about BP regulation. In this study, we take a different approach, making use of recent advances in single cell sequencing and study expression of genes at BPs in individual cells to learn about their regulation. Recent developments in single cell genomics allow the measurement of RNA expression in individual cells with a similar accuracy as compared to bulk-sequencing of RNAs [15, 16]. This advance has been used to define previously overlooked cell types and expression heterogeneity, *e.g.*, [17].

We used novel and previously produced single cell RNA-seq (scRNA-seq) data for HepG2 and K562 cells to investigate the expression behavior of genes at a BP. We found that four reproducible expression categories exist in BPs and that in the majority of the cases, one gene at a BP shows much higher expression than the other one. Using high resolution histone modification datasets produced at IHEC standards [18] by the DEEP consortium or made available by ENCODE [19], we find novel associations of different structural and epigenetic features in these categories.

## 2 Results

To understand the regulation of the two genes at a bidirectional promoter, we propose a novel approach to exploit RNA-seq data at the single cell level, in contrary to the existing studies that rely on bulk RNA-seq data. Bulk RNA-seq masks gene expression across individual cells, and thus may hide interesting expression patterns of bidirectional gene pairs (Figure 1A).

**Figure 1.**
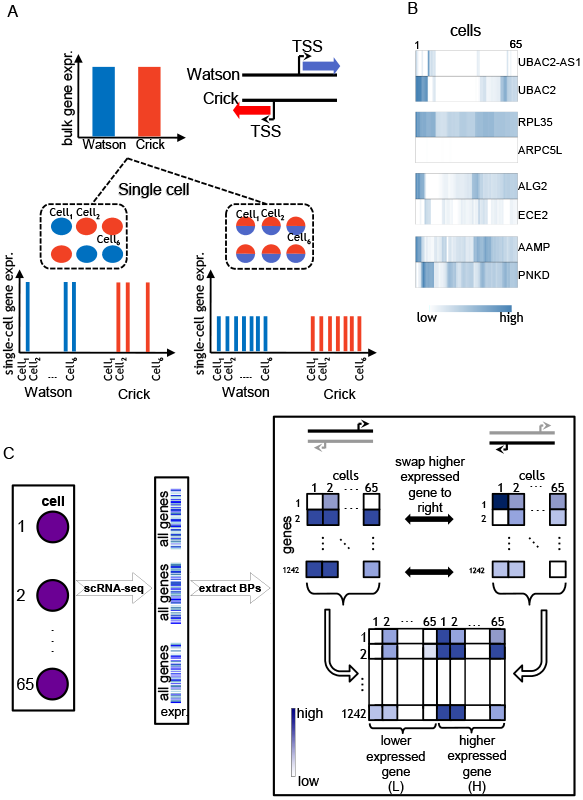
Advantages of studying BPs at single cell level. A) An illustration of a BP, defined based on two genes located on opposing strands of DNA (Watson and Crick). Bulk RNA measurements at the BP may hide complexity of BP gene regulation. This is shown in the left single cell expression scenario, where one of the genes is expressed and the other is silent in the same cell compared to the other scenario where single cell expression agrees with bulk measurements. B) Heatmaps of 65 single cell RNA-seq expression measured in four bidirectional promoters (TPM, HepG2 cells). C) After single cell sequencing and estimating the gene expression of all genes in a cell, a set of 1,242 BPs was extracted. Single cell expression of either genes of a BP was arranged in two separate matrices for which the rows represent the BPs and columns the cells. Next, we swap the higher expressed gene to the matrix on the right and lower expressed one to the left. The resulting matrices are combined into one joint BP single cell expression matrix.

Figure 1B illustrates examples of single cell expression patterns in HepG2 cells for selected BPs. It can be noted that, for instance, the magnitude of expression of the *ALG2*, *ECE2* gene pair alternates across the cells, meaning that in some cells *ALG2* is higher expressed than *ECE2* and vice versa. Similarly, *AAMP* and *PNKD* genes exhibit this alternation, but more frequently. These observations motivated us to inspect such diversities in a systematic manner by forming an expression matrix specific to BPs for clustering analysis.

### 2.1 Four states of transcription with distinct bidirectional characteristics

We form an individual matrix of all BPs representing the single cell expression of the gene located on the Watson strand (Watson matrix). Similarly, we construct the same matrix for the gene on the Crick strand (Crick matrix) (Figure 1C). To simplify the follow-up analyses, we swap a row of the Watson matrix with the corresponding Crick row, if the average single cell expression of the former is lower than the latter. In this way, for a given BP, we always keep the higher expressed gene (H) on the right side and the lower expressed one (L) on the left. Next, we form the final swapped BP matrix, where the rows represent the bidirectional genes (N=1,242) and the columns represent the cells (twice the number of single cells); the first half of the columns represent cells’ expression of L genes and the second half represent the same for H genes. Since, the combined matrix contains the joint expression information for both genes of a BP in each row, we used hierarchical clustering to group the BPs according to their similarity in single cell expression patterns. This led to four distinct transcription states in both cell lines (Figure 2A HepG2, and Supplementary Figure 2A K562) with the following characteristics: 1) *Bidirectional Lowly Expressed* (*BLE*), where both genes of a BP are lowly expressed, 2) *Bidirectional Weak Difference* (*BWD*), where the H gene is higher expressed than the L gene with a weak difference between the two, 3) *Bidirectional Strong Difference* (*BSD*), where the H gene is much higher expressed than the L gene and higher than in *BWD*, 4) *Bidirectional No Difference* (*BND*), where both genes of a BP are expressed relatively at the same rate.

**Figure 2.**
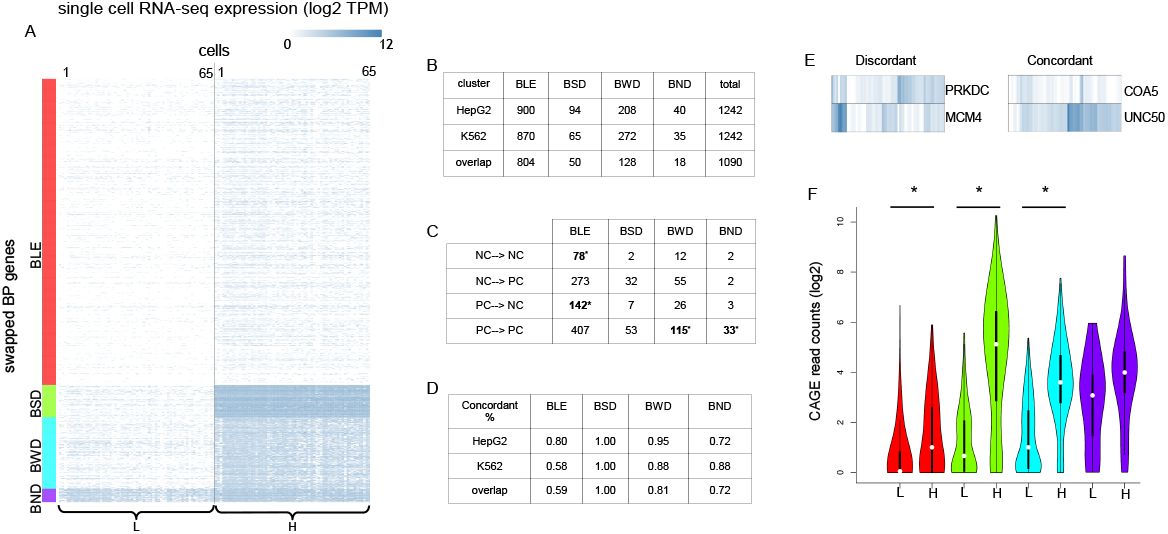
Single cell RNA-seq expression in bidirectional promoters. A) Hierarchical clustering of the HepG2 single cell transcript expression matrix visualized as a heatmap (log2, TPM). The four distinct clusters (*BLE*, *BSD*, *BWD*, *BND*) are referred to as *transcription state* in this manuscript. B) Number of BPs falling into each transcription state in HepG2 and K562 cells and their overlap. C) Number of BPs falling into the gene product categories (NC→ NC, NC→ PC, etc.) in HepG2. Statistically enriched values are shown in bold (Hypergeometric test p<0.05). D) Ratio of *concordant* BPs shown separately in each state for both cell lines as well as their overlap. E) Examples of *concordant* and *discordant* BPs in HepG2. F) CAGE read counts, measured for each bidirectional gene (L and H), shown for each transcription state. Color code as in A. Significant differences are marked with * (paired and two-sided Mann-Whitney test, *p* <0.05).

The data regarding the frequency and type of BPs in each state is provided in figures 2B,C. Figure 2B reveals that most of the BPs associated to these states are common between the two cell lines (1,090 out of 1,242). We investigated whether the transcription state was related to the type of genes encoded in a BP. We found that for both cell lines the *BWD* and *BND* states are enriched with BPs (hyper-geometric test, p ≤ 0.05), where both bidirectional genes are annotated as protein-coding (*PC* → *PC*, Figure 2C, Supplementary Figure 2B). On the other hand, the *BLE* state is enriched with BPs of either two non-coding genes (*NC* → *NC*) or where the L gene is annotated as protein-coding and the H gene as non-coding (*PC* → *NC*).

The single cell data allowed us to estimate the frequency of (*concordant* or *discordant*) gene signatures of BPs in all states for both cell lines (2D,E). The *BLE* state was overall lowly expressed and due to stochasticity of expression, it is difficult to find a consistent pattern for this particular state. On the other hand, *BSD* state consists of BPs where one gene’s expression is always higher than the other, thus we obtained a *concordant* ratio of 1. As expected, the *BND* state is showing some of the smallest *concordant* ratios, i.e, highest *discordance*, which points to the frequent alternations (switches) occurring in the expression of the genes in this state.

Figure 2F illustrates that the CAGE expression distributions follow the characteristics attributed to each cluster (similarly for the bulk RNA-seq and CAGE in K562 cell line, Supplementary Figure 2C,D). However, it is worth mentioning that performing the clustering based on the bulk data, either RNA-seq or CAGE did not lead to a reproduction of the transcription clusters based on single cell RNA-seq, due to measuring a population of cells in bulk assays (data not shown).

The representation used in Figure 1D is concise, but it does not provide a suitable visualization to explore the associations between L and H genes in the same cell. Therefore, to quantitatively assess the relation between single cell expression of bidirectional genes in these states, we computed, for each BP, the correlation between expression of L and H genes across single cells (Figure 3A, data shown for both cell lines). The correlation analysis showed that the *BND* state has the highest correlation. On the contrary, the *BSD* state revealed lower correlation, which suggests a more independent regulation of its bidirectional genes.

**Figure 3.**
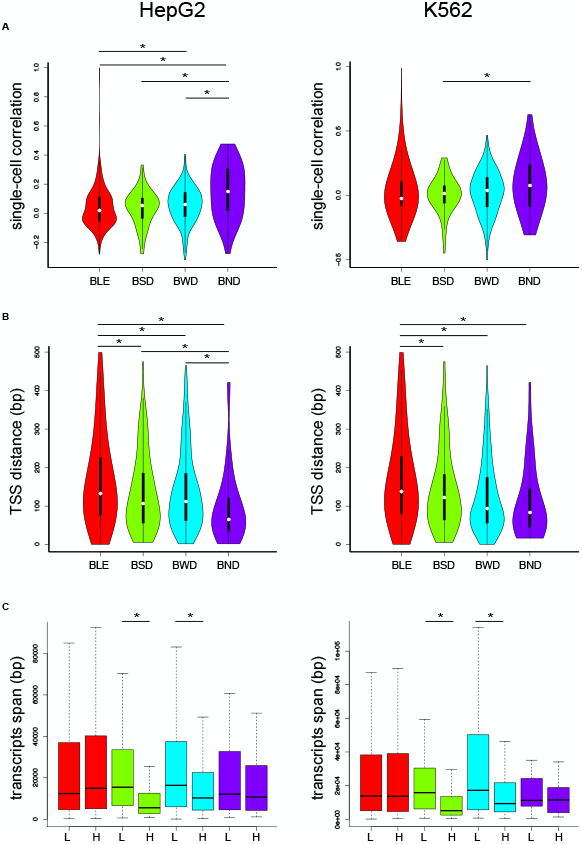
Structural features of BPs for HepG2 (left column) and K562 cells (right column). A) Distributions of Pearson correlation coefficients (y-axis) calculated from all single cell measurements for each BP in one of the states (x-axis). B) Distributions of TSS distance of BPs in each state. C) Length distributions of *transcripts span* for L and H genes of BPs shown in each state. Significant differences are marked with a * (paired and two-sided Mann-Whitney test, *p* <0.05). For all subfigures the color-coding is consistent with Figure 1D.

To address which mechanism(s) are involved in driving such differences in regulation of BPs, we explored the following aspects: 1) structural features, 2) epigenetic signals, and 3) transcriptional regulatory elements.

#### Structural features associated with transcription states

We first tested whether the distance between TSSs of bidirectional genes was associated with the transcription states. Figure 3B depicts the distributions of TSS distances in each state for both cell lines. We observed that the *BLE* state exhibits significantly larger TSS distances compared to the other states (t-test, p ≤ 0.001). On the contrary, the *BND* state had the smallest median distance (significant for HepG2, t-test p ≤ 0.05). This observation together with the correlation analysis in Figure 3A suggests that the smaller distance may influence recruitment of a common regulatory complex that facilitates the simultaneous regulation of both genes.

As the scRNA-seq protocol measures steady-state fully elongated mRNAs, we wondered whether the length of the transcribed region differs in the genes associated to the BPs. For this, we examined the region spanned by all transcripts originating from transcription start sites within 2 kb from the most 5^′^ TSS of a BP gene, a region we refer to as *transcripts span* (see Materials and methods). Surprisingly, this length was significantly smaller (Mann-Whitney test, p-value ≤ 0.05) for the H genes of states *BSD* and *BWD* compared to their counterpart L genes. Connecting this observation to the actual transcription expression depicted in Figure 1D for these two states suggests that the expressions of L and H genes are inversely related to their *transcripts spans* in BPs. To elucidate whether this association holds for all genes or only BPs, we measured the *transcripts span* for all 63,678 annotated genes in the human genome. We found no association of *transcripts span* with gene expression for all genes (Supplementary Figure 2F), indicating that such structural configuration might be specific to BPs. As the estimated TPM values are derived from the exonic regions only, we further examined the transcript length by measuring the exonic region of all transcripts initiating within the 2 kb from the most 5′ TSS of a BP gene (Supplementary Figure 2H,I). We found a slight increase of TPM values for the larger genes, regardless of considering all genes or only BP (Supplementary Figure 2F).

We also investigated whether the difference in GC-content could be involved in driving variation on the observed expression patterns, but we found no apparent differences (Supplementary Figure 2G).

#### Histone modification and DNaseI patterns reflect the characteristics observed in transcription states

To explore the role of epigenetics in transcription states observed in Figure 1D, we produced seven histone modifications (H3K4me1, H3K4me3, H3K36me3, H3K27me3, H3K9me3, H3K27ac, and H3K122ac) and DNaseI-seq data for HepG2 cells within the DEEP consortium. Further we obtained data for DNaseI-seq and all modifications, except H3K122ac, for K562 cells from [19]. Figure 4 depicts the normalized read counts measured around the TSSs of bidirectional genes stratified according to the transcription states for all HepG2 datasets (similarly, for K562 in Supplementary Figure 3A). Generally, we observed that the epigenetics data show specific patterns related to these states. For instance, it is notable that the *BLE* state had the lowest abundance for HMs associated with active promoters (H3K4me1/3, H3K36me3, H3K27ac, and H3K122ac) and highest for H3K27me3 and H3K9me3 that are mostly associated with repressed promoters [20]. On the other hand, the *BND* state exhibited the very opposite behavior to *BLE*, reflecting their expression characteristics observed in Figure 1D.

**Figure 4.**
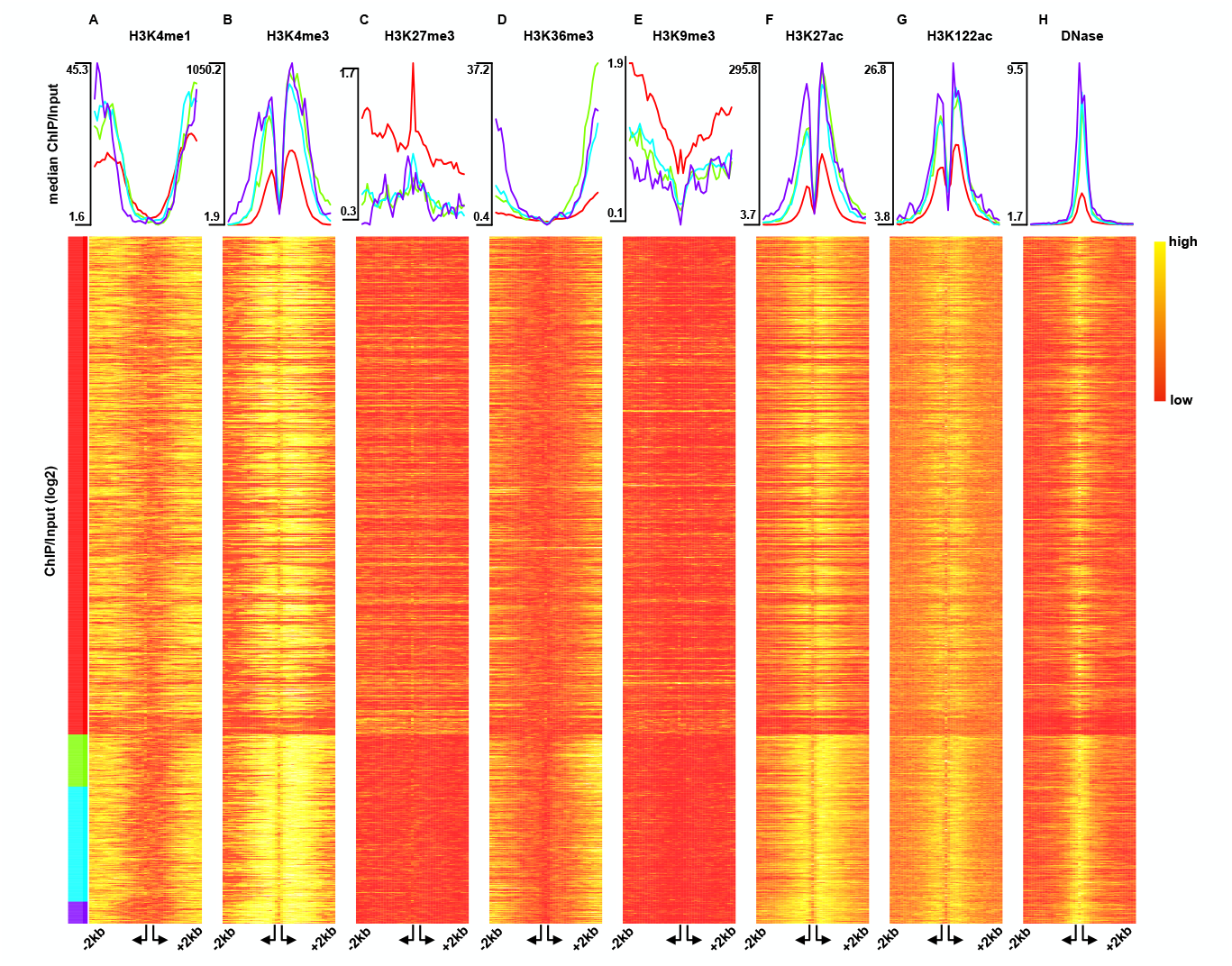
Epigenetic characteristics in transcription states in HepG2 cells. A-G) Histone modification (ChIP/Input) shown as median profiles (top panel) and log-transformed values as heatmap (bottom panel). H) DNase1-seq median profiles (top panel) and log-transformed raw counts (bottom panel). Arrangement of genes as in Figure 1D. The reads are measured in 40 bins of size 100 bp forming a window of size 4000 bp centered around the TSSs, with an additional variable bin between the TSSs.

Another interesting observation is the agreement of the elongation mark profiles, H3K36me3, with the *transcripts span* distribution shown in Figure 3C. In general, the larger the increase of the H3K36me3 mark the shorter the *transcripts span* for the gene. For instance, the *BSD* state that has the shortest *transcripts span* exhibits the sharpest increase in its H3K36me3 profile downstream of the H gene’s TSS. This fits to the previous observation that the H3k36me3 mark increases gradually and peaks at the end of genes [21] and we can observe that general trend for the *transcripts span* on our data as well (Supplementary Figure 3B).

The DNaseI-seq profile of the *BND* state revealed not only the highest signal, but also the widest spread around the TSS compared to the other states. This agrees with the observation that there is similar amount of single cell transcription for both genes.

Due to recent reports about small promoter-associated RNAs [22, 23], we obtained small RNA data [19] for HepG2 and K562 samples (see Methods) and grouped them according to the defined transcription states. Although we observed residual small RNA expression in the vicinity of the bidirectional TSSs, we found no consistent patterns associated with the transcription states (Supplementary Figure 3C).

We also examined the average methylation profiles obtained in the four transcription states (Supplementary Materials and Methods) due to the previously reported associations with gene expression [24, 25]. The results were consistent with other studies where higher level of DNA methylation coincided mostly with silent genes (*BLE*). Consistent with the enrichment of HMs, genes in the *BND* state showed the least amount of DNA methylation (Supplementary Figure 3D).

#### The BND state coincides with strongest regulatory activity

It was shown that certain TFs preferentially bind to bidirectional promoters [13, 14]. As we observed that the DNA accessibility profiles differed among the transcription states (Figure 4H), we were encouraged to investigate binding of transcription factors. We obtained ChIP-seq data for several transcription factors [19] (44 for HepG2 and 50 for K562). One hypothesis was that there may exist TFs that bind in the proximal region of a BP and influence gene expression as was observed in our transcription states.

To test this, we defined a novel enrichment score tailored to BPs (Supplementary Figure 4A), which preserves the spatial distribution of the ChIP-seq signal along a BP. We applied the enrichment analysis for both cell lines (HepG2 in Figure 5A and K562 in Supplementary Figure 4B). As expected, states with higher expression showed more TF binding in general. However, we could not pinpoint distinct TF subsets that associate with only one of the states. Instead, the states *BSD*, *BWD* and *BND* showed enrichment for many of the TFs that we analyzed. We wondered whether the number of TFs that are regulating a BP differed in those states. Figures 5B,C represent the number of positively enriched TFs per BP for each state in both cell lines. The *BND* state showed the highest percentage of positively enriched TFs (t-test, p ≤ 0.05) suggesting that more TFs are required to regulate gene expression in this state.

**Figure 5.**
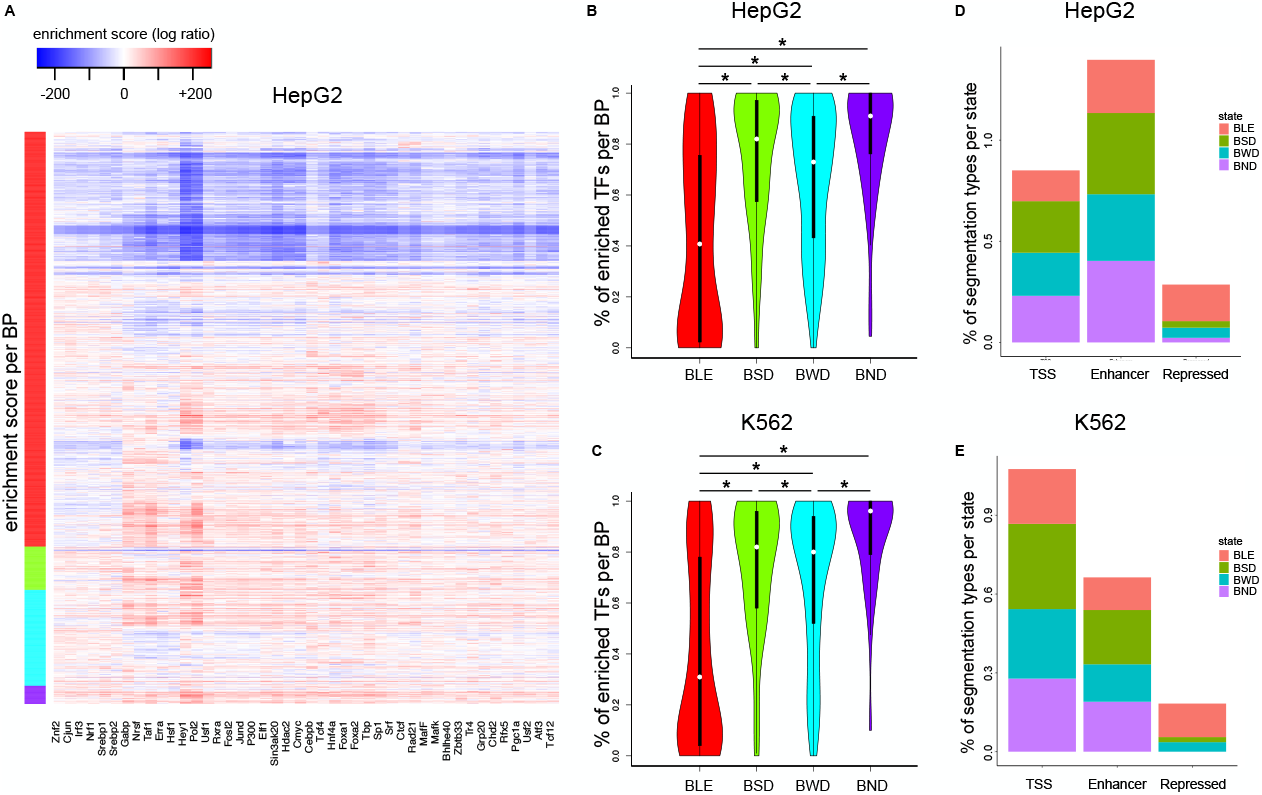
Transcriptional regulatory features in the transcription states. A) Heatmap of TF enrichment scores (log ratio against background) for each BP (row) in HepG2 cells. BPs are sorted as in Figure 1D. B,C) Distributions of percentages of TFs per BP (enrichment score in A > 0) in each state for HepG2 (top panel) and K562 (bottom panel). D,E) ChromHMM annotations, summarized into the types: *TSS*, *Enhancer*, and *Repressed*, are shown as percentages in a bar plot per state (see Materials and methods).

Next, we tested whether specific genomic regions, such as enhancers, are associated with these four transcription states. For this, we inspected the genome-wide segmentation of HepG2 and K562 cells using an 18-state ChromHMM model [26] (Supplementary Figure 5, Supplementary Materials and Methods). For simplification we collapsed all TSS-related, enhancer-related, and repression-related ChromHMM states into *TSS*, *Enhancer*, and *Repressed*, respectively. We assigned all the remaining chromatin states to *Others* (data not shown). The results provided in Figures 5D,E suggest that the enhancer-related regions are the most frequent amongst the *BSD* and *BND* states, reflecting their stronger expression profiles. In the case of HepG2 (Figure 5D), this quantity is even higher than the number of *TSS* regions. Concurrent with Figure 4 most of the repressed regions belong to the *BLE* state, where genes were lowly expressed.

## 3 Discussion and conclusions

In this work we compared single cell expression of genes at BPs. Currently, we only have access to single cell protocols for RNA-seq, and other techniques for quantification of transcription start sites cannot be used [6, 4, 27]. Thus, other effects on the mRNA steady state level, *e.g.* post-transcriptional regulation, may influence the gene clustering produced. Here, we have used two high quality single cell datasets for ENCODE cell lines allowing us to benefit from a plethora of epigenomic datasets, which are available or have been produced in this work. We found that 88% of the BPs have the same transcription state in scRNA-seq data despite the difference in origin of HepG2 and K562 cells, which suggests that the majority of these configurations may be stable for many cell types.

In previous work that has analyzed BP regulation, analyses were often limited to a certain configuration at the BP, *e.g.* a non-coding gene upstream of a coding gene, therefore care has to be taken when comparing to previous studies. Here, we have limited our results to annotated protein- or non-coding genes that originate from a bidirectional promoter. We found that the BPs that show similar expression for both genes are mostly restricted to a configuration with two protein-coding genes. It was shown previously that core promoter strength differs for genes with bidirectional expression and unidirectional promoters [5]. Here, we show that, beyond differences in the strength of the core promoter, the number of TF regulators that bind to BPs with high bidirectional expression is largest compared to all other expression configurations we observed. In this analysis we used several ChIP-seq datasets for TFs and developed a BP-specific enrichment analysis approach that measures spatial differences in read coverage along the BP regions compared to the median background, unified in a single quantity for each BP and TF. This is different to other studies that have compared TF ChIP-seq data at BPs, e.g. [14], where the background often were unidirectional promoters rather than all BPs. Thus, to find enrichment in the observed states we properly adjust for the fact that there are two genes, which are regulated by TF binding.

We observed that the *BND* state shows the largest (although not strong) single cell correlation values and that there is a trend with correlation at BP genes being inversely proportional to TSS distance (Figures 2A,B). A similar observation was recently made for BPs in the rice genome with correlation measured over several bulk RNA-seq datasets [28]. Small distance between the two TSSs may ease the coupled regulation of transcription from both, for example through a shared or co-regulated Mediator complex [29].

We also found that the *transcripts span*, the genomic region covered by all transcripts that start in the vicinity of the TSS, was imbalanced for the *BSD* and *BWD* states, with the shorter span linked to the highly expressed gene at the BPs. One possibility is that shorter regions of elongation lead to faster transition cycles for Pol2, assuming similar elongation rate of both genes at a BP. This could be a mechanism by the cell to create imbalanced expression output from a shared regulatory region of two BP genes. Anecdotally, we investigated bulk GRO-cap data for K562 cells [4], and found that the amount of transcription initiation is more similar for both genes at a BP in our states (Supplementary Figure 2E), compared to the amount of stable RNAs expressed (CAGE and RNA-seq). Even though the initiation is the same we get significantly different steady-state transcript expression, which could be explained by the difference in length of the genomic region to be elongated, here referred to as *transcripts span*. Once single cell measurements of nascent transcription are available, one could investigate the difference in elongation and transcriptional initiation in these BPs.

**Figure 6.**
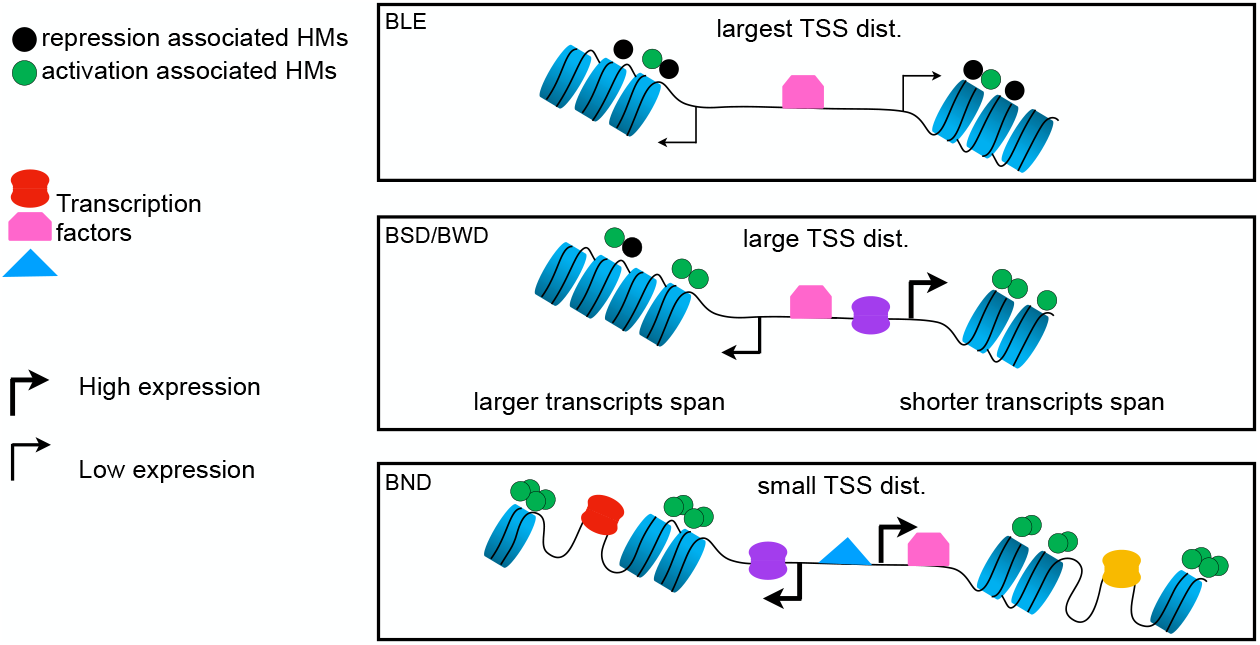
Hypothetical model of three different genomic architectures underlying epigenetic regulations of BPs. BPs that drive single cell expression patterns observed in the *BLE* state show large TSS distance and higher abundance of repression associated marks and depletion of most TFs. *BSD* and *BWD*, on the other hand, exhibit smaller TSS distance and more TF binding compared to *BLE*. In addition, the *transcripts span* of the H gene is observed to be significantly smaller compared to the L gene. BPs categorized in *BND* show the smallest TSS distance with the most TF binding events that require more accessible DNA to regulate both the L and H genes.

Taken together, we observed three different genomic and epigenomic architectures underlying single cell transcription states in BPs. We propose a model depicted in Figure 5 to describe these architectures. This model supports distinct characteristics of the *BLE* state, where the bidirectional genes were lowly expressed. They mostly exhibited large TSS distance and more prevalence of repression associated HMs, fewer regions of accessible DNA, and less TF binding. The *BSD* and *BWD* states, on the other hand, had reduced TSS distance in comparison with *BLE* and more abundance of activation associated HMs as well as higher rate of TF binding. Interestingly, the *transcripts span* associated to the H gene of BPs in these states was observed to be shorter than the L one. Lastly, *BND* showed strongest single cell co-expression and smallest TSS distance among the states. Furthermore, we observed the widest accessible regions of DNA, the largest number of binding TFs and highest amount of activation related HMs.

Although the transcription state definition was based on the single cell data, several bulk datasets showed consistent and matching patterns for those states. Our results suggest that novel statistical methods can be developed to deconvolute masked subpopulations of cells measured with different bulk epigenomic assays with the help of single cell RNA-seq data. Further advances in single cell sequencing technologies [30] are necessary such that we can measure not only RNA expression, but also TF binding and histone modifications in single cells to understand the hidden complexity, in particular, in BP regulation.

## 4 Methods

### Datasets and pre-processing

#### Single cell RNA-seq

Single HepG2 cells were manually picked to prepare poly-A enriched cDNA libraries using Smart-seq2 as described by [31] with modifications. Briefly, 65 single cell samples were supplemented with 0.5 *μl* of a 1:40,000 dilution of the Ambion ERCC RNA Spike-In Mix 1 (Thermo Sientific, #4456740). After cell lysis polyadenylated mRNA was reverse transcribed using a biotinylated template switch oligo (5^′^-Biotin-AAGCAGTGGTATCAACGCAGAGTACATrGrG+G-3^′^) with two riboguanosines (rG) and one LNA-modified guanosine (+G) at the 3^′^ end. Preamplified cDNA (18 PCR cycles) was purified with Agencourt Ampure XP Beads (Beckman Coulter, #A 63881) in a 1:1 ratio. cDNA quality of 8 random samples was assessed on the Agilent 2100 Bioanalyzer (Agilent Technologies, #G2938C) using the Agilent high-sensitivity DNA kit (Agilent Technologies, # 5067- 4626). Sequencing libraries were prepared using the Nextera XT DNA Sample Preparation Kit (Illumina, #FC-131- 1024) with approximately 480 *pg* of cDNA in a 4 *μl* tagmentation reaction followed by a dual indexing PCR with 9 cycles. Individual single cell libraries were pooled and purified with 0.8 X Agencourt Ampure XP Beads. The library pool was sequenced on a HiSeq 2500 (Illumina) using the TruSeq SBS Kit v3-HS (Illumina, #FC-401- 3001) in a single read run with 90 bp read length.

#### Single cell transcript expression

The TPM values for transcript isoforms of each Ensembl gene (GRCh37) were computed using RSEM [32]. To attribute the transcription expression to each bidirectional gene, we summed the isoform TPM values of transcripts that had their annotated TSS within a 2 kb window downstream of the most 5^′^ TSS of that gene.

#### HepG2 and K562 datasets

Epigenomic data for the HepG2 cell lines have been produced by the DEEP consortium and are deposited at the European Genome-Phenome Archive under the accession number EGAS00001001656. The rest of the data, K562 (HM-ChIP-Seq, TF-ChIP-seq, CAGE), and HepG2 (TF-ChIP-seq, CAGE) were obtained from the ENCODE portal.

### Bidirectional promoter (BP) gene set

The BP dataset contained 1,242 divergent promoters with two core promoter elements, obtained from annotated ENSEMBL genes (GRCh37.75), such that the distance between TSSs of each BP does not exceed 500 bp. This set excludes loci overlapped by any other annotated gene region (±2 kb from the TSS).

### Clustering BPs into four states

Hierarchical clustering (TPM values) using the complete linkage method with Euclidean distance as distance metric was applied on the swapped BP matrix using R.

### Constructing the single cell TPM matrix for BPs

For a particular *BP*, *BP_i_* = (*g_crick,i_, g_watson,i_*), we compute the sum of TPM values across single cells as following:

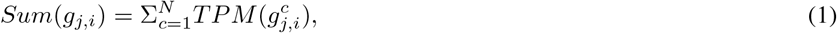
where *N* denotes the number of single cells, and *TPM* 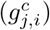 returns the TPM value for gene *j* ∈ {*crick, watson*} of *BP_i_* in cell *c*. The orientation of genes at a BP is not specific to the DNA strand, but the lower expressed gene of a BP is always swapped to the left and higher expressed gene to the right. In this way, without loss of generality, all analyses correctly adjust for differences of expression. Precisely, we define *g_H,i_* denoting the gene of *BP_i_* having higher expression as follows:

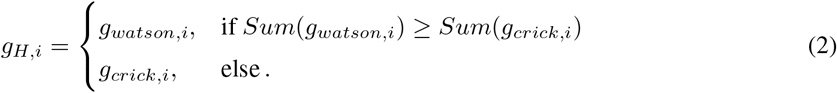

Similarly, we define *g_L,i_* denoting the gene of *BP_i_* having lower expression:

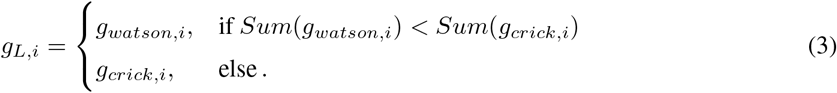

After defining *g_H,._* and *g_L,._* for each BP, we form the single cell matrix for BPs, *scBP*, as follows:

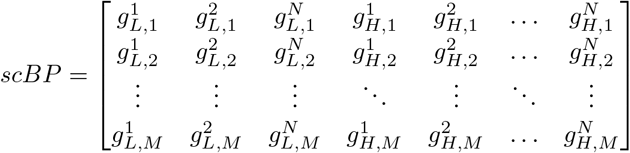

### Imputation of drop-outs

To address the bias caused by drop-outs, we performed the most recent and accurate drop-out imputation tool called scImpute [33], which aims to improve the single-cell data quality by removing the effects of drop-outs without introducing new biases to the data. scImpute has two parameters. *K* denotes the number of existing cell types in the data, which we set to 1, as we work on the cell lines. The second parameter *t* controls the drop-out probabilities. The authors show that their results are robust to different parameter values, therefore, we carried on with the default of 0.5 for this parameter. The comparison between raw and imputed read counts performed on the bidirectional genes is shown in Supp. Fig. 1A for both HepG2 and K562. The Pearson correlation between imputed and raw data in both cell lines is ~ 1.

#### Quality of scRNA-seq

Imputed expression of bidirectional genes averaged over single cells was compared with their corresponding bulk RNA-seq expression. For both, HepG2 and K562, the single cell expression agrees well with bulk measurements (Spearman correlation coefficient of ~ 0.8, Supplementary Figure 1B). Additionally, the imputed TPM values were divided into three intervals, 1 < *TPM* < 10, 10 ≤ *TPM* ≤ 100, *TPM* > 100 to account for the number of genes falling in those intervals per cell (Supplementary Figure 1C, and similarly for the imputed read counts in 1D).

### Bidirectional gene signature: *concordant* or *discordant*

We define two types of signatures to address the changes in bidirectional gene expression. Intuitively, if the two genes are mostly expressed in a consistent manner across the single cells, for example one is always higher than the other, this would be considered as *concordant* signature. However, if the expression of these two genes flips across cells, we refer to this case as *discordant*. To analytically differentiate between both signatures for each pair of genes in a BP, we performed the Wilcoxon signed rank test on their imputed single cell expression (BPs where both genes had zero expression in all cells were removed for the test). If the p-value after using Benjamini-Hochberg multiple testing correction is smaller than or equal to 0.05, the gene pair is considered to be *concordant*. The number of *concordant* BPs normalized by the total number of BPs in a given cluster is defined as *concordant* ratio.

### Enrichment of gene products categorized according to transcription states

We categorized the gene product annotations into two groups, protein-coding (*PC*) and the rest as non-coding (*NC*). In the context of BPs, we introduce a new notation, *gp* ∈{*NC* → *NC, NC* → *PC, PC* → *NC, PC* → *PC*}, representing the gene products of a pair of genes. We measured the occurrences of each of the above four categories for the gene pairs of our transcription states as shown in Figures 1F and Supplementary Figure 2C. To compute the enrichment of such occurrences, we applied a hypergeometric test on their contingency table, *C* ∈ ℤ^4×4^, where *C_i,j_* represents the frequency of the *j^th^* gene product category in the *i^th^* state. Precisely, let *h*(*x*; *N, n, k*) be the hypergeometric distribution, where *N* denotes the population size, *n* denotes the sample size, *k* is the frequency of successes in the population, and *x* represents the frequency of successes in the sample. To apply this distribution to each entry *C_i,j_* of the contingency matrix *C*, we used the following setup:

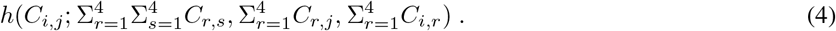

The p-value derived from this test was used to quantify the significance of enrichment of a gene product category in a particular state.

### Enrichment of TF ChIP-seq data

To preserve the spatial distribution of the TF ChIP-seq signal around the promoter, the ChIP-seq reads are counted in bins of size 100 bp forming a window starting at the TSS of each bidirectional gene and extending up to 2000 bp downstream of each of two TSSs (Supplementary Figure 4A). An additional bin with variable size is allocated to count the reads falling within the region between the TSSs of the two bidirectional genes. The 20 bins from the *L* gene, the bin for region between both TSSs, and the 20 bins from the *H* gene are all combined into one vector of size 41 that represent the binned ChIP-seq signal per BP for a particular TF. To compute the enrichment score of the *i^th^* TF at a particular BP, we define:

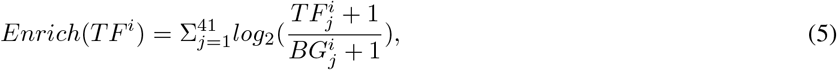
where *TF^i^* is the signal measured for *i^th^* TF (for HepG2, *i* ∈ {1, …, 44} and for K562, *i* ∈ {1, …, 50}) at the given BP. 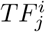 denotes the read counts measured at the *j^th^* bin of *TF^i^* signal and 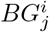 denotes the median of *TF^i^* signal measured at the *j^th^* bin across all BPs.

### Definition of transcribed regions

For each gene, we consider all the annotated transcripts that start within 2 kb downstream of the most 5^*1*^ TSS of the gene. We measured the length of the exonic region encompassed by these transcripts, which we refer to as transcript length. Similarly, the exonic and intronic region spanned by those transcripts is referred to as *transcripts span*. For instance, if the following transcripts start downstream within 2 kb of the most 5^′^ TSS, *T*_1_ = (*start* : 0, *end* : 1000), *T*_2_ = (*start* : 200, *end* : 3000), *T*_3_ = (*start* : 200, *end* : 2000), then the *transcripts span* would be equal to (*start* : 0, *end* : 3000), where *start* and *end* are relative coordinates to the most 5^′^ TSS. Note that all regions in this interval are considered, regardless of their exonic or intronic annotations. Also note that other transcripts of the gene that would start outside of the 2 kb region are not considered for the definition of *transcripts span* or transcript length.

## 5 Declarations

### Ethics approval and consent to participate

Not applicable.

### Consent for publication

Not applicable.

### Availability of data and material

All sequencing DEEP data used for this study have been deposited at the European Genome-Phenome Archive under the accession number EGAS00001001656. The source code for the conducted analyses and code for making the the figures can be found in the GitHub repository viahttps://github.com/SchulzLab/scBP.

### Author’s contributions

FBA was involved in all computational analysis supported by KN and JF; KK and SF generated the single cell RNA-seq data; NG, GG and KN generated and processed Dnase1-seq and DNA methylation data; KK generated ChIP-seq data; MB, AS and PR generated and processed bulk RNA-seq data; BF, GZ, KN, PE, BH, BB, JE performed mapping and management of sequencing data; PE generated ChromHMM segmentation; MHS designed and coordinated the study supported by FBA, NG, JW, GG, TL, TM. MHS and FBA wrote the manuscript with contributions from other authors.

## Acknowledgements

We thank Alex A Pollen and the SRA archive for help with obtaining the single cell data for K562. We are thankful to all labs that contributed to the ENCODE data used.

